# On distinguishing between canonical tRNA genes and tRNA gene fragments in prokaryotes

**DOI:** 10.1101/2022.07.05.498093

**Authors:** Peter T.S. van der Gulik, Martijn Egas, Ken Kraaijeveld, Nina Dombrowski, Astrid T. Groot, Anja Spang, Wouter D. Hoff, Jenna Gallie

**Affiliations:** Centrum Wiskunde & Informatica, Amsterdam, 1090 GB, The Netherlands; Department of Evolutionary and Population Biology, Institute for Biodiversity and Ecosystem Dynamics, University of Amsterdam, Amsterdam, 1090 GE, The Netherlands; Leiden Centre for Applied Bioscience, University of Applied Sciences Leiden, Leiden, 2333 CR, The Netherlands; Department of Marine Microbiology and Biogeochemistry, NIOZ, Royal Netherlands Institute for Sea Research, Den Burg, 1790 AB, The Netherlands; Department of Microbiology and Molecular Genetics, Oklahoma State University, Stillwater, Oklahoma, 74078, USA; Department of Evolutionary Theory, Max Planck Institute for Evolutionary Biology, Plön, 24306, Germany

## Abstract

Automated genome annotation is essential for extracting biological information from sequence data. The identification and annotation of tRNA genes is frequently performed by the software package tRNAscan-SE, the output of which is listed for selected genomes in the Genomic tRNA database (GtRNAdb). Here, we highlight a pervasive error in prokaryotic tRNA gene sets on GtRNAdb: the mis-categorization of partial, non-canonical tRNA genes as standard, canonical tRNA genes. Firstly, we demonstrate the issue using the tRNA gene sets of 20 organisms from the archaeal taxon Thermococcaceae. According to GtRNAdb, these organisms collectively deviate from the expected set of tRNA genes in 15 instances, including the listing of eleven putative canonical tRNA genes. However, after detailed manual annotation, only one of these eleven remains; the others are either partial, non-canonical tRNA genes resulting from the integration of genetic elements or CRISPR-Cas activity (seven instances), or attributable to ambiguities in input sequences (three instances). Secondly, we show that similar examples of the mis-categorization of predicted tRNA sequences occur throughout the prokaryotic sections of GtRNAdb. While both canonical and non-canonical prokaryotic tRNA gene sequences identified by tRNAscan-SE are biologically interesting, the challenge of reliably distinguishing between them remains. We recommend employing a combination of (i) screening input sequences for the genetic elements typically associated with non-canonical tRNA genes, and ambiguities, (ii) activating the tRNAscan-SE automated pseudogene detection function, and (iii) scrutinizing predicted tRNA genes with low isotype scores. These measures greatly reduce manual annotation efforts, and lead to improved prokaryotic tRNA gene set predictions.

## INTRODUCTION

Canonical transfer RNAs (tRNAs) implement the genetic code by coupling mRNA codons to their respective amino acids during protein synthesis. Of the 64 codons in the genetic code, three generally serve as stop codons. The remaining 61 (‘sense’) codons are decoded into 20 amino acids by tRNAs with various complementary anticodons. Given that some anticodons can recognize more than one codon – through wobble base pairing ^1,2^ – organisms carry fewer than 61 types of tRNA ^3^. Together with knowledge of wobble base pairing rules, systematic studies of tRNA gene sets have yielded important generalities regarding tRNA requirements for efficient translation; standard tRNA gene sets have been identified for each of the three domains of life ^4,5^. In Archaea – the focus of this work – a standard set of 46 tRNA types, each encoded by a single gene copy, has been proposed (examples are depicted in Figure 1A). Standard tRNA gene sets are poised to play a pivotal role in understanding tRNA gene set function and evolution. However, a range of reported deviations from these sets (*e*.*g*., ^3,6^) may be masking their role in tRNA biology. Specifically, two types of tRNA gene set deviations can occur: (i) evolutionary adaptations of the translationally active tRNA gene set in some organisms (see below), and (ii) artifactual deviations due to imperfections in genome sequencing/assembly, or downstream computational analyses of putative canonical tRNA genes. This manuscript aims to distinguish between these two cases, using the tRNA gene sets carried by the archaeal Thermococcaceae family as an example.

**Figure 1.**
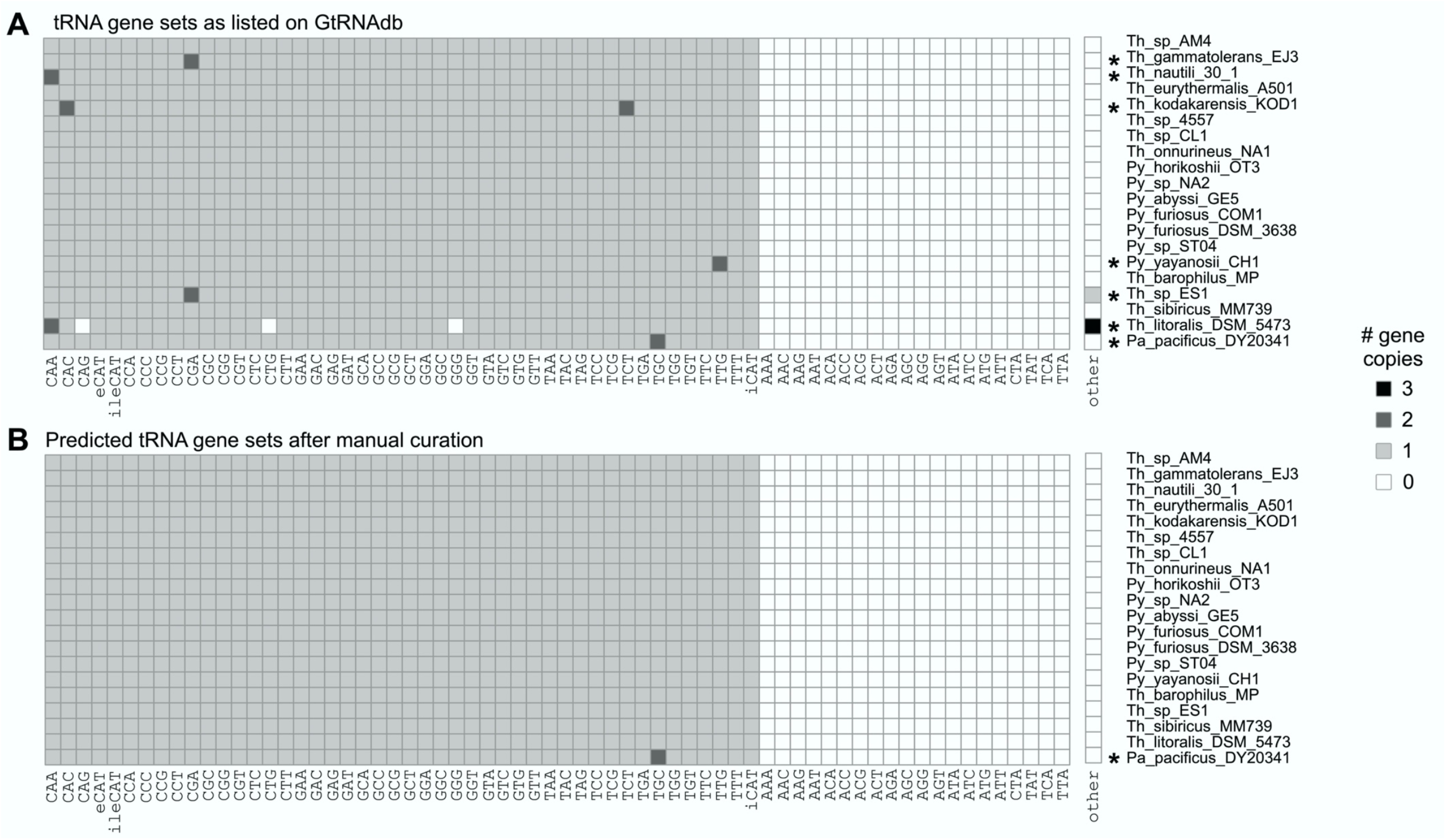
Heatmap depicting putative tRNA gene sets for 20 Thermococcaceae family organisms. (**A**) tRNA gene sets as listed on GtRNAdb (predicted via tRNAscan-SE 2.0.6, with pseudogene detection deactivated), and (**B**) following the manual annotation procedure described in the Materials and Methods. Organisms with non-standard tRNA gene sets are marked by an asterisk (seven in panel A, one in panel B).

As a result of remarkable progress in experimental methods for determining the genome sequences, automated annotation has become an essential tool for extracting biological information from sequence data. In the field of tRNA biology, several tRNA gene prediction tools are available ^7–13^, the most widely used of which is tRNAscan-SE ^7,14,15^. tRNAscan-SE effectively performs the challenging tasks of identifying putative tRNA genes from input DNA sequence, and annotating each predicted gene – including the definition of the anticodon and amino acid carried (*i*.*e*., the tRNA isotype). Additionally, tRNAscan-SE distinguishes between three isotypes carrying the anticodon 5’-CAU-3’: (i) initiator tRNA-Met(CAU), which serves in the initiation stage of translation, (ii) elongator tRNA-Met(CAU), which functions in peptide chain elongation, and (iii) tRNA-Ile(CAU), in which the anticodon is post-transcriptionally modified to recognize isoleucine codon 5’-AUA-3’ ^16^.

The final output of tRNAscan-SE is a biochemically detailed, automatically annotated list of putative tRNA genes from the input DNA sequence (*e*.*g*., genome assembly, plasmid). The tRNAscan-SE output for a selected set of complete (or near-complete) genomes is collected in the publicly accessible Genomic tRNA database (GtRNAdb), with all predicted tRNA genes falling into one of five categories (‘tRNAs decoding standard 20 AA’, ‘Selenocysteine tRNAs (TCA)’, ‘Possible suppressor tRNAs (CTA,TTA,TCA)’, ‘tRNAs with undetermined or unknown isotypes’, and ‘Predicted pseudogenes’) ^3,17^. The accumulation and comparison of complete canonical tRNA gene set data (*i*.*e*., tRNA genes in the first category) paves the way for conclusions regarding the evolution of tRNA gene sets across the tree of life ^5,18–26^. In addition, such information may have important medical implications ^27–31^. For example, both prokaryotic and eukaryotic tRNA gene sets have been shown to evolve rapidly in response to changing translational demands ^32–34^, and a correlation has been demonstrated between tRNA gene copy number and bacterial growth rate ^35^. However, conclusions regarding tRNA gene sets are dependent on the accuracy and proper use of the tRNA detection software.

Here, we evaluate the performance of tRNAscan-SE, through a detailed manual inspection of the tRNA genes it identifies and annotates, within a set of 20 genomes from the archaeal family Thermococcaceae. This family is an archaeal clade within the phylum *Euryarchaeota* (according to the <u>N</u>ational <u>C</u>enter for <u>B</u>iotechnology Information (NCBI) taxonomy ^36^), or the phylum *Methanobacteriota* (according to the <u>G</u>enome <u>T</u>axonomy <u>D</u>ata<u>b</u>ase, GTDB ^37^). The Thermococcaceae family consists of at least six genera (*i*.*e*., *Pyrococcus, Thermococcus, Thermococcus A, Thermococcus B, Thermococcus C*, and *Palaeococcus*) that are typically found in hydrothermal marine environments (^38^, chapter 27). Members of the Thermococcaceae family reportedly adhere very closely to the standard tRNA gene set of Archaea (^38^, page 364). Given this information, the proportion of Thermococcaceae genomes listed in GtRNAdb as encoding variant tRNA gene sets is – as described below – unexpectedly high (seven deviant genomes out of the 20 listed genomes = 35 %). However, manual annotation of the predicted tRNA gene sets resolves many of the apparent differences, leaving only a single probably truly divergent tRNA gene set.

## MATERIALS AND METHODS

### 20 Thermococcaceae tRNA gene sets

All Thermococcaceae family tRNA gene sets available in GtRNAdb (Data Release 19; June 2021) ^3^ were downloaded (see Supplementary Table S6). Details of the 20 organisms – including members of the *Pyrococcus, Thermococcus, Thermococcus A, Thermococcus B*, and *Palaeococcus* genera – are provided in Supplementary Table S1. In addition, the tRNA gene sets of these organisms were predicted from the genome assemblies (downloaded from NCBI) using locally run tRNAscan-SE (version 2.0.6, the same version that was reportedly used for the current GtRNAdb listings for all 20 organisms), with standard settings for Archaea (option -A) and pseudogene detection activated (see Supplementary Table S7). To display the output, options -H and --detail were added.

### Manual annotation of tRNA gene sequences

Putative tRNA genes of interest were manually annotated using the following six steps: (i) Identify the 3’ CCA terminus (nucleotides 74-76), the discriminator nucleotide (nucleotide 73) and the acceptor stem (nucleotides 1-7, base pairing with nucleotides 72-66) ^39^. (ii) Working 3’ to 5’, identify the T-arm prior to nucleotide 66 (>>>>>UUCRANN<<<<<, where R is base A or G, and each >< indicates a base pair). (iii) Identify the section between the T-arm and the anticodon-arm (nucleotides 44-45-46-47-48 or 44-45-47-48, or an extra arm). (iv) Identify the anticodon-arm (>>>>>NNNNNNN<<<<<). With very few exceptions, the anticodon loop consists of precisely seven nucleotides (N), and very often begins with CU. (v) Identify the D-arm (>>>xNNNNNNNNx<<<, where x denotes a degree of flexibility in the number of Ns). (vi) Identify nucleotides 8-9 (between the acceptor stem and D-arm) and 26 (between the D-arm and anticodon arm), which play a role in realization of the three dimensional structure of tRNA molecules.

### Phylogenetic tree construction

The Thermococcaceae family phylogenetic tree in Supplementary Figure S1 was generated by first identifying suitable marker genes in genomes from 20 *Thermococcus* and three *Methanococcaceae* (outgroup) genomes, as described previously ^40^. Briefly, HMM profiles of an extended TIGRFAM database were queried against a protein database of all 23 archaeal genomes, using hmmsearch v3.1b2 and the following settings: hmmsearch --tblout sequence_results.txt -o results_all.txt --domtblout domain_results.txt --notextw extended_TIGR.hmm All_Genomes.faa. The extended TIGRFAM database can be found at https://zenodo.org/record/3839790#.YjByaVzMI3g. Only hits with an e-value <= 1e-3 were included, and the best hit per protein based on the bit score and e-value was selected. Markers with =>10% of duplicated proteins were excluded, leaving 45 markers for downstream analyses. Note that TIGR00335 and TIGR00483 were excluded from these analyses, because they comprised a large number of paralogues. The final set of 43 marker proteins (listed in Supplementary Table S5) was extracted from the larger protein database and individually aligned using MAFFT LINS-i v7.407 (settings: –reorder) ^41^ and trimmed using BMGE v1.12 (settings: -t AA -m BLOSUM30 -h 0.55) ^42^. Single-protein phylogenies were inferred with IQ-TREE v2.1.2 (settings: -m LG -T AUTO -keep-ident --threads-max 2 -bb 1000 -bnni) ^43^. Trees were manually inspected and all potential paralogues removed. The remaining proteins were realigned, trimmed (as above) and concatenated using a custom perl script; catfasta2phyml.pl (https://github.com/nylander/catfasta2phyml). The phylogenetic tree was inferred based on a final alignment of 12,403 positions using IQ-TREE (v2.1.2, -m LG+C60+F+R -T AUTO --threads-max 80 -bb 1000 -alrt 1000) ^43^, visualised using FigTree (v1.4.4) and annotated using Adobe Illustrator CC2018.

### 210 archaeal tRNA gene sets

The tRNA gene sets of 217 Archaea are currently listed in GtRNAdb (Data Release 19; June 2021). For seven of these organisms, the underlying genome assemblies were no longer available from NCBI (*Aciduliprofundum* sp. MAR08-339, *Methanococcus maripaludis* C6, *Methanococcus maripaludis* C7, *Methanococcus voltae* A3, *Methanolinea tarda* NOBI-1, *Methanosarcina barkeri* str. Fusaro, and *Saccharolobus solfataricus* 98/2 (formerly *Sulfolobus solfataricus* 98/2)). The remaining 210 organisms span five phyla and include 59 Crenarchaeota, 138 Euryarchaeota, 1 Korarchaeota, 1 Nanoarchaeota, and 11 Thaumarchaeota. The genomes of these organisms are available on Zenodo (https://zenodo.org/record/6782366#.YsQS5pDP01B). For each of these 210 organisms, the tRNA gene sets and associated scores listed in GtRNAdb were downloaded (resulting in the ∼10,000 tRNA genes listed in Supplementary Table S7). In addition, the genome assemblies (including chromosomes and plasmids) used to generate the GtRNAdb entries were downloaded from NCBI, and tRNAscan-SE (version 2.0.6) was locally run on each genome, using standard settings for Archaea (options -A, -H, --detail) and with pseudogene detection activated. In accordance with expectations, ∼10,000 tRNA genes were predicted (Supplementary Table S7).

## RESULTS

### GtRNAdb lists divergent tRNA gene sets for seven of 20 Thermococcaceae genomes

The tRNA gene sets of 20 genomes belonging to the Thermococcaceae family are listed in GtRNAdb (Data Release 19; June 2021) (Supplementary Table S1). Our phylogenetic tree of these 20 genomes confirmed the phylogeny-informed taxonomy of the Thermococcaceae family established by Rinke *et al* ^44^ and accurately resolved the relationships between the five genera to which these genomes belong, with *Palaeococcus* forming the earliest diverging branch (Supplementary Figure S1).

All 20 of the genomes of interest are complete genomes assembled into single circular contigs, with an average size of 1.9 Mb (ranging from 1.7 Mb to 2.2 Mb) and a mean genomic GC content of 47 % (ranging from 40 % to 56 %). Genome descriptions have been published for 18 of the 20 genomes (see Supplementary Table S1). Inspection of the associated GtRNAdb tRNA gene set entries revealed that, of the 20 genomes, only 13 (65 %) are predicted to carry the standard archaeal tRNA gene set of 46 canonical tRNA types encoded by single-copy genes (see Introduction), while the remaining seven (35 %) are predicted to differ. There are 15 distinct anomalies across the seven deviant genomes, consisting of twelve additional and three absent putative tRNA genes (Table 1; Figure 1A).

**Table 1.** Fifteen instances in which the 20 Thermococcaceae genomes listed in GtRNAdb reportedly differ from the standard archaeal tRNA gene set.

| No.<sup>a</sup> | Affected isotype(s)<sup>b</sup> | Size (bp)<sup>c</sup> | Isotype Model Score<sup>d</sup> | Organism | Locus (strand) | tRNA scan-SE<sup>e</sup> | GtRNA db<sup>f</sup> | Cat.<sup>g</sup> |
| --- | --- | --- | --- | --- | --- | --- | --- | --- |
| 1 | (+)Ser-CGA | 53 | 10.4 | *T. gammatolerans* EJ3 | 621,672 - 621,724 (-) | Ps | St | 1b |
| 2 | (+)Leu-CAA | 218 | 2.3 | *T. nautili* 30-1 | 706,451 - 706,668 (+) | Ps | St | 1a |
| 3 | (+)Arg-TCT | 83 | 21.4 | *T. kodakarensis* KOD1 | 320,075 - 320,157 (-) | Ps | St | 1b |
| 4 | (+)Val-CAC | 72 | 83.4 | *T. kodakarensis* KOD1 | 62,858 - 62,929 (-) | St | St | 1b |
| 5 | (+)Gln-TTG | 90 | 13.4 | *P. yayanosii* CH1 | 1,321,660 - 1,321,749 (-) | Ps | St | 1b |
| 6 | (+)Ser-CGA | 69 | 30.9 | *T. sp.* ES1 | 747,982 - 748,050 (+) | Ps | St | 1b |
| 7 | (+)Und-NNN | 127 | 16.8 | *T. sp.* ES1 | 603,860 - 603,986 (-) | Und | Und | 1b |
| 8 | (+)Leu-CAA | 76 | 6.3 | *T. litoralis* DSM 5473 | 326,949 - 327,024 (+) | Ps | St | 1b |
| 9-10 | (+)Leu-NAG<br>(-)Leu-CAG | 88 | 135.1<br>(134.9) | *T. litoralis* DSM 5473 | 648,439 - 648,526 (+) | Ps | St | 2 |
| 11-12 | (+)Pro-NGG<br>(-)Pro-GGG | 78 | 120.3<br>(120.3) | *T. litoralis* DSM 5473 | 546,230 - 546,307 (-) | Ps | St | 2 |
| 13-14 | (+)Und-NTG<br>(-)Gln-CTG | 76 | 117.9<br>(117.9) | *T. litoralis* DSM 5473 | 825,204 - 825,279 (-) | Ps | St | 2 |
| 15 | (+)Ala-TGC | 77 | 122.6 | *Pa. pacificus* DY20341 | 1,527,238 - 1,527,314 (-) | Ps | St | 3 |

Rows are ordered according to the phylogenetic tree in Supplementary Figure S1.
<sup>a</sup>Anomaly number (1-15)
<sup>b</sup>(+) indicates additional tRNA genes, (-) indicates missing tRNA genes. Pairs of additional and missing tRNA genes are placed together (see also Supplementary Text S1).
<sup>c</sup>Length in base pairs of the predicted tRNA gene
<sup>d</sup>Isotype score reported by GtRNAdb
<sup>e,f</sup>Classification by locally run tRNAscan-SE (version 2.0.6; pseudogene detection active), or GtRNAdb (Ps=Predicted pseudogenes, St=tRNAs decoding standard 20 AA, Und=tRNAs with undetermined or unknown isotypes)
<sup>g</sup>Category that the anomaly falls into. 1=erroneously predicted tRNA gene resulting from (1a) CRISPR-Cas activity or (1b) integration of a mobile genetic element (*i.e.*, *attR* sites), 2=isotype misassignment, 3=legitimate additional tRNA gene).

When the same set of genomes were analyzed by running tRNAscan-SE locally (version 2.0.6, see Materials and Methods), six of the genes causing tRNA gene set deviations were flagged as possible pseudogenes. Inspection of the documentation for the entries of these genomes on GtRNAdb indicated that automated pseudogene detection was not activated during the associated tRNAscan-SE run, resulting in the inclusion of the six potential pseudogenes within the canonical, standard tRNA gene sets for the organisms involved.

### Manual annotation reveals only a single divergent tRNA gene set

Given the unexpectedly high degree of deviation from the standard archaeal tRNA gene set, the 929 tRNA genes predicted across the 20 Thermococcaceae genomes were subjected to careful manual inspection (see Materials and Methods). This revealed that the 15 observed anomalies fall into three categories, each of which is described below (see also Table 1).

#### Category 1: Erroneously predicted tRNA genes arising from tRNA gene fragments

Eight of the 15 observed anomalies result from the erroneous listing of partial tRNA gene sequences as either standard tRNA genes (seven cases) or genes encoding tRNAs of undetermined isotypes (one case) in GtRNAdb (Table 1). As described further below, we found that the partial tRNA sequences are the result of two processes: (i) the acquisition of fragments of tRNA genes from viral genomes by CRISPR-Cas activity (accounting for one anomaly; category 1a in Table 1), and (ii) the partial duplication of tRNA genes during the integration of horizontally transferred genomic elements (accounting for seven anomalies; category 1b in Table 1).

*Thermococcus nautili* 30-1 (NCBI Reference Sequence NZ_CP007264.1) is predicted to carry two distinct genes encoding Leu-CAA (positions 10,012–10,099 and 706,451–706,668). The first of these has characteristics typical of canonical leucyl-tRNA genes (*e*.*g*., 88-bp long, isotype score 134.8), while the second does not (*e*.*g*., 218-bp long, very low isotype score of 2.3). Furthermore, the RNA product of the second is predicted to form an unusual secondary structure, in which the T-arm is the only readily identifiable tRNA feature (see Supplementary Text S1). Closer inspection of the *T. nautili* 30-1 genome reveals that the second, atypical Leu-CAA sequence occurs within a CRISPR array of over 40 identical short repeats interspersed with unique spacer sequences that are derived from foreign, invading DNA (Figure 2A) (for a recent review of CRISPR-Cas systems, see ^45^). The tenth spacer of this array appears to be derived from a viral tRNA gene, encompassing the T-arm and one side of the acceptor stem (Figure 2B-C). A search for nucleotide sequence matches to the spacer-10 sequence and viral DNA sequences (using the Nucleotide Basic Local Alignment Search Tool, blastn) revealed a good match between the spacer and the Thr-TGT gene of several isolates from the *Myoviridae* family, a family of bacteriophage known to infect Archaea and Bacteria (Figure 2D). Based on these results, we conclude that the tRNA gene fragment in *T. nautili* 30-1 is derived from a viral tRNA gene. While tRNA genes have been documented in a range of bacteriophage genomes ^46,47^, CRISPR spacers do not commonly contain viral tRNA gene fragments ^48,49^; to the best of our knowledge, this is the first report of a tRNA gene fragment appearing in a CRISPR-Cas array. Together, these results are consistent with the CRISPR-Cas-mediated incorporation of a viral tRNA gene fragment into the *T. nautili* 30-1 genome, which is erroneously listed as a canonical, standard tRNA gene in GtRNAdb.

**Figure 2.**
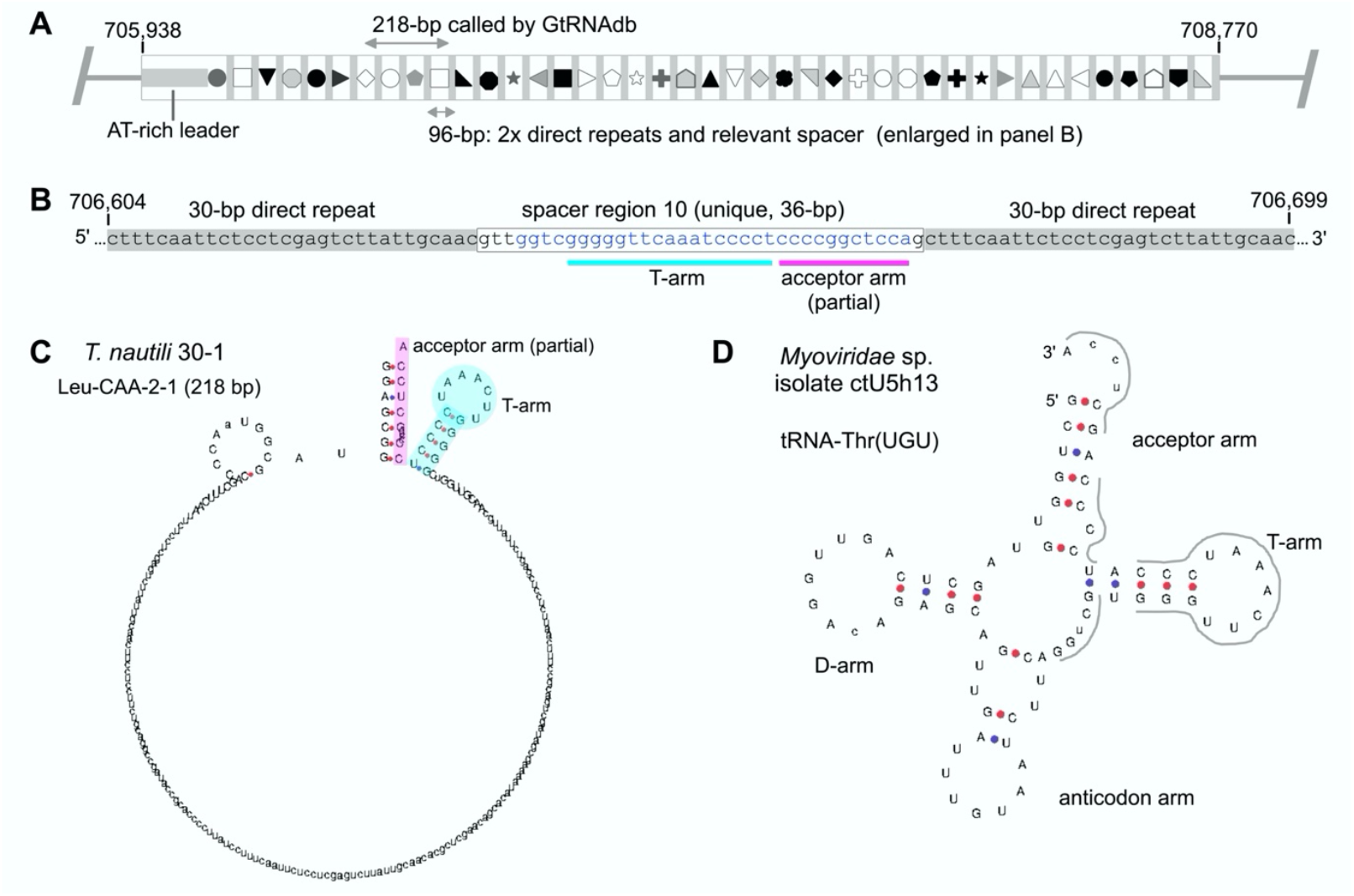
A partial tRNA gene resulting from CRISPR-Cas activity in *T. nautili* 30-1 (category 1a in Table 1). (**A**) Cartoon depicting the *T. nautili* 30-1 CRISPR-Cas locus of interest (genomic bases 705,938–708,770). The array consists of 41 identical 30-bp direct repeats (grey rectangles) interspersed with 41 unique spacers of ∼36 bp (various shapes). The 218-bp Leu-CAA-2-1 tRNA identified by GtRNAdb contains sequence from spacers 7-10 and repeats 7-9 (bases 706,451– 706,668). Of these, spacer 10 contains the actual tRNA gene remnant. (**B**) Enlargement of spacer 10 and flanking repeats (bases 706,604–706,699). Blue text indicates the 32-bp region matching to viral tRNA sequence (see panel D). (**C**) The partial tRNA gene sequence in spacer 10 is predicted by tRNAscan-SE to form a secondary structure in which the T-arm (turquoise) and part of the acceptor stem (magenta) are identifiable. (**D**) The T-arm and acceptor stem of tRNA-Thr(UGU) from *Myoviridae* isolates (*Myoviridae* sp. Isolate ctU5h13 ^70^ is shown here) are a good match for spacer 10; the sequence matches 29 of the 32 blue bases in panel B (matches indicated by thick grey lines).

Inspection of a further seven of the deviating tRNA genes revealed that they involve fragments of tRNA genes resulting from insertion of mobile genetic elements into the genome. A particularly clear example of such an element has been previously reported in *Pyrococcus yayanosii* CH1 (NCBI Reference Sequence NC_015680.1; ^50^), one of the 20 Thermococcaceae genomes: the ∼21.4 kb mobile genetic element PYG1 lies between partial and complete Gln-TTG copies (Figure 3A-C) ^51^. Both the partial and complete Gln-TTG sequences are listed in GtRNAdb as distinct, standard tRNA genes (Figure 3D-E). Our dataset includes a further six examples of partial tRNA genes resulting from similar integration events, with evidence for the integration of between ∼9 kb and ∼23 kb of viral DNA into various archaeal tRNA genes (see Supplementary Table S2). These findings are in line with published results showing that the incorporation of mobile genetic elements (*e*.*g*., temperate phages, integrative and <u>c</u>onjugative <u>e</u>lements (ICEs)) into prokaryotic genomes is a widely occurring source of tRNA gene fragments; tRNA genes are common targets for the integration of such elements, presumably as a result of tRNA gene stability and pervasiveness ^52–54^. Indeed, manual inspection of arbitrary prokaryotic GtRNAdb entries rapidly revealed numerous instances of similar partial tRNA genes in other Archaea, and Bacteria, many of which are erroneously listed as standard, canonical tRNA genes on GtRNAdb (for examples from various taxa, see Supplementary Table S3).

**Figure 3.**
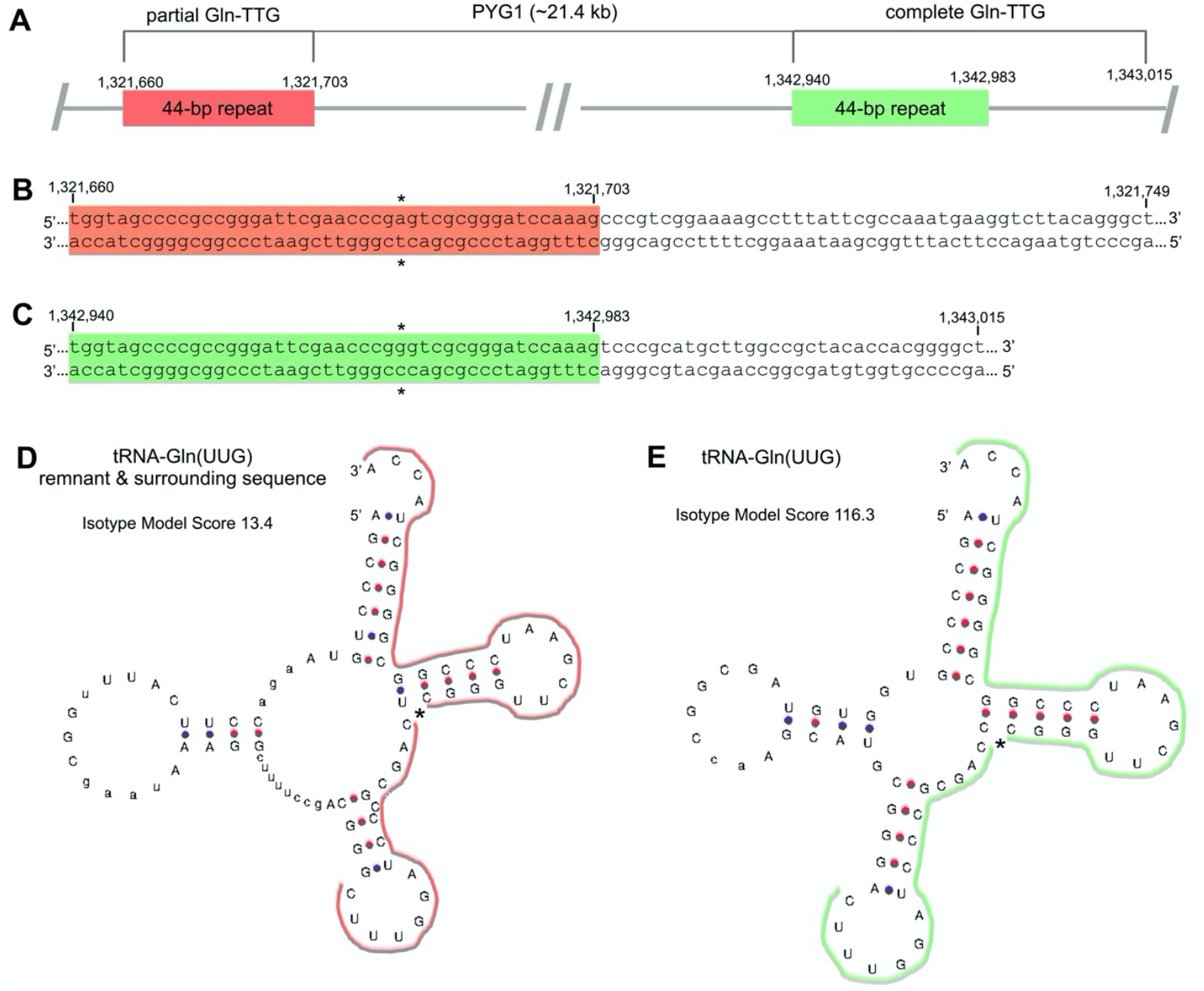
A partial tRNA gene resulting from integration of a mobile genetic element in *P. yayanosii* CH1 (category 1b in Table 1). (**A**) *P. yayanosii* CH1 contains a partial and a complete copy of the Gln-TTG tRNA gene, separated by the ∼21.4 kb genomic island, PYG1 ^51^. PYG1 is flanked by two 44-bp direct repeats (1,321,660–1,321,703 in red; 1,342,940–1,342,983 in green), only the second of which forms part of a standard, canonical Gln-TTG gene. (**B, C**) Primary sequences of the 90-bp and 76-bp Gln-TTG genes listed as putative standard tRNA genes in GtRNAdb. The 44-bp direct repeats are highlighted (red/green), with * indicating a 1-bp difference. (**D, E**) tRNAscan-SE-derived secondary structures of the 90-bp and 76-bp Gln-TTG gene sequences, respectively. Thick red and green lines indicate the 44-bp repeats.

The ICE integration process typically results in one complete and one partial copy of the target tRNA gene at opposing ends of the integrated element, providing a reliable method for their delineation ^55^. Notably, the identification of these types of partial tRNA genes, and their full tRNA gene counterparts, can also provide evidence of genome rearrangement events. For example, our analysis indicates the integration of a ∼20.7 kb prophage into the canonical Arg-TCT gene of the common ancestor of *Thermococcus gammatolerans* EJ3 and *Thermococcus kodakarensis* KOD1, followed by an inversion event in the KOD1 lineage that led to the occurrence of the partial and full tRNA gene copies on opposite strands of the chromosome, and a considerable distance apart (∼179 kb; see Supplementary Table S2).

#### Category 2: Erroneously assigned tRNA isotypes resulting from ambiguous input sequence

The *Thermococcus litoralis* DSM 5473 genome (NCBI Reference Sequence NC_022084.1; ^56^) contains seven of the 15 anomalies, including all three of the missing tRNA genes (Leu-CAG, Pro-GGG, Gln-CTG) and four additional tRNA genes (Leu-NAG, Pro-NGG, Und-NTG, Leu-CAA) (Figure 1A; Table 1). Careful analysis of the primary and secondary structures strongly indicates that three of the additional tRNA genes account for the three missing tRNA genes: additional gene Leu-NAG is the apparently missing gene Leu-CAG, Pro-NGG is Pro-GGG, and Und-NTG is Gln-CTG (see Supplementary Text S1 for tRNA structures). In each case, the presence of one or more N (*i*.*e*., identity unknown) nucleotides in the tRNA gene sequence – and the *T. litoralis* DSM 5473 genome sequence – hampers the ability of tRNAscan-SE to predict the anticodon and/or isoacceptor family of the putative tRNA gene. Together, these errors account for a further six of the anomalies (Table 1). The seventh anomaly is described above in category 1b (a tRNA gene fragment resulting from the integration of incoming DNA). Hence, after manual annotation, the *T. litoralis* DSM 5473 genome is predicted to encode the standard archaeal tRNA gene set.

#### Category 3: Legitimate, additional tRNA genes

In the two categories above, manual annotation has accounted for 14 of the 15 observed tRNA gene set anomalies. The remaining anomaly is an additional copy of the Ala-TGC gene in *Palaeococcus pacificus* DY20341 (NCBI Reference Sequence NZ_CP006019.1; ^57^). The two Ala-TGC gene copies reportedly occur within distinct, distally separated ribosomal RNA (*rrn*) operons, each consisting of a 16S gene, Ala-TGC, and a 23S gene. With the exception of an additional 32-bp stretch of DNA in one of the 23S genes, the two reported *rrn* loci are identical in primary sequence (including Ala-TGC). Notably, the other 19 Thermococcaceae genomes each encodes only a single *rrn* operon (and hence only one Ala-TGC gene; see Supplementary Table S4) ^58^. These results are consistent with intra-genomic duplication of the *rrn* operon in *Pa. pacificus* DY20341, leading to an actually divergent tRNA gene set in this taxonomically distinct organism (see Supplementary Figure S1). However, it should be noted that long stretches of nearly identical DNA sequence (such as *rrn* operons) pose considerable technical challenges for genome sequencing and assembly, and hence that *rrn* copy numbers (and any tRNA genes contained) may be inaccurate ^59,60^.

The results so far show that, after manual annotation, the tRNA gene sets encoded in the 20 Thermococcaceae genomes adhere closely to the standard archaeal tRNA gene set of 46 single-copy tRNA genes; only a single probable deviation – duplication of Ala-TGC in *Pa. pacificus* DY20341 – remains (Figure 1B).

### The tRNAscan-SE isotype score as an indicator of non-canonical tRNA genes

Manual annotation requires expert knowledge of tRNA structure, and detailed manual annotation may be too time consuming for very large data sets. It is therefore desirable to devise automated methods for distinguishing between canonical and non-canonical tRNA genes. As reported above, the automated detection of possible pseudogenes offered by tRNAscan-SE is valuable in this regard, as it flags multiple tRNA gene fragments identified here as probable non-canonical tRNA genes. In addition, during our manual annotation process, we noted lower than usual isotype scores for non-canonical tRNA genes. Specifically, nine predicted tRNA genes contributing to the deviations of the standard archaeal tRNA gene set exhibited relatively low isotype model scores (see Table 1). Therefore, we performed a more systematic analysis of the value of both approaches (pseudogene detection feature and isotype score) for detecting non-canonical tRNA genes. To this end, we obtained the isotype scores of all predicted tRNA genes for the 20 Thermococcaceae genomes from GtRNAdb and locally run tRNAscan-SE (version 2.0.6; see Materials and Methods), and extracted the tRNA genes flagged as possible pseudogenes from the output of locally run tRNAscan-SE.

In each case, 929 tRNA genes were predicted, with isotype scores ranging between 2.3 and 138.2 (median=115.8; Figure 4A-B; Supplementary Table S6). While the isotype scores of most predicted tRNA genes fell into a Gaussian-like large peak with a maximum of ∼120, eight putative tRNA genes – each corresponding to a partial tRNA gene sequence listed in Table 1 – received low isotype scores (<85; Figure 4C-D). One of these low-scoring putative tRNA genes – *Thermococcus* sp. ES1 Und-NNN (127-bp long; isotype score 16.8) – is called as a non-canonical tRNA of undetermined or unknown isotype by tRNAscan-SE (and in GtRNAdb). A further six were identified as pseudogenes by locally run tRNAscan-SE, yet listed as standard tRNA genes in GtRNAdb. The lack of pseudogene definition in GtRNAdb appears to result from the active disablement of pseudogene checking by tRNAscan-SE (as can be seen under the “Run Options/Stats” tab for individual organisms listed in GtRNAdb). The decision to deactivate the possible pseudogene detection feature when scanning archaeal (and bacterial) genomes for the current GtRNAdb listings was made in order to avoid mis-classifying some low-scoring, but likely canonical, putative tRNA genes as pseudogenes (Todd Lowe, personal communication).

**Figure 4.**
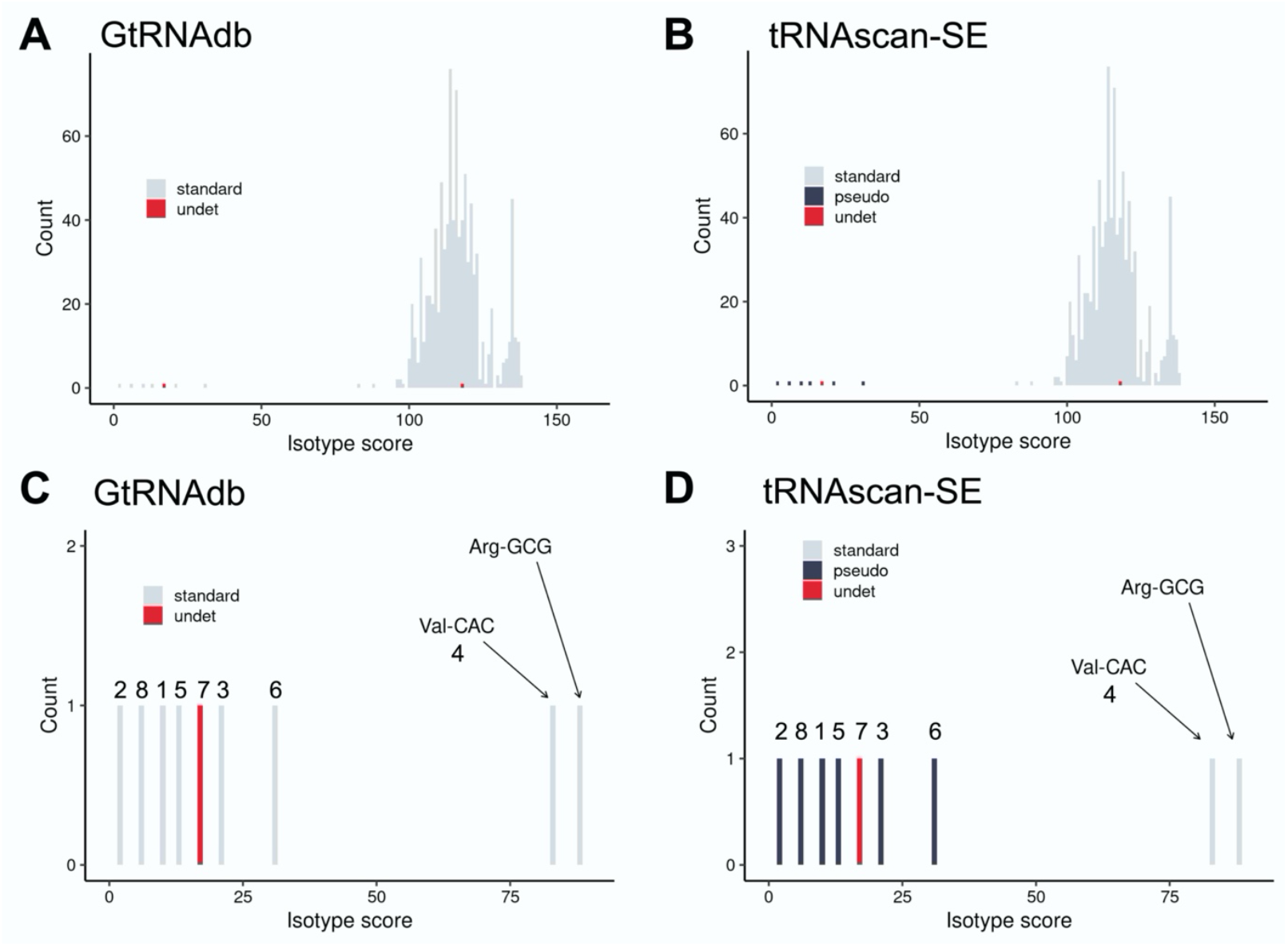
tRNAscan-SE isotype score as a predictor of canonical versus non-canonical tRNA genes in the 20 Thermococcaceae genomes. Histograms of the frequency of tRNA isotype scores from all 929 tRNA genes listed in (**A**) GtRNAdb, and (**B**) predicted by locally run tRNAscan-SE, for the 20 Thermococcaceae genomes. (**C-D**) Closeup of isotype scores 0-100 in panels A and B. Numbers 1-8 relate to the anomalies listed in Table 1. tRNA gene categories are generated by tRNAscan-SE. Note that in panels A and C (GtRNAdb) the pseudogene category (‘pseudo’) is absent, due to disabling of pseudogene detection function (see ‘Run Options/Stats’ tab in GtRNAdb listings).

The final relatively low-scoring putative tRNA gene – Val-CAC of *T. kodakarensis* KOD1 (isotype score 83.4) – is called as a standard tRNA gene by both tRNAscan-SE and GtRNAdb. However, manual inspection of the primary sequence and the predicted secondary structure reveals defects in the acceptor stem and D-arm that render the putative tRNA highly unlikely to be canonically functional (see Supplementary Text S1). The tRNA with the next lowest isotype model score was *Pyrococcus furiosus* DSM 3638 Arg-GCG (isotype score 88.1). Given that this single-copy tRNA gene encodes an essential tRNA isotype, it is highly likely to give rise to a functional, canonical tRNA. Indeed, its relatively low isotype score can be accounted for by the absence of a single nucleotide in the acceptor stem (G70), which could arise from a point mutation or an error in the DSM 3638 reference genome sequence (see Supplementary Text S1). Overall, these results show that for the tRNA gene dataset from the 20 Thermococcaceae genomes, a combination of (i) the automated detection of probable pseudogenes/fragments, (ii) the tRNAscan-SE isotype score, and (iii) manual inspection of tRNA genes with borderline isotype scores (∼60-90) can be used as a valuable indicator of the nature of putative tRNA genes (canonical versus non-canonical).

To investigate whether the above observations extend to a wider archaeal dataset, the distribution of isotype scores for all putative tRNA genes in 210 complete (or nearly complete) archaeal genomes listed in GtRNAdb were examined. For these 210 organisms, approximately 10,000 putative tRNA genes were listed in GtRNAdb and also predicted by locally run tRNAscan-SE (the latter with pseudogene detection activated; see Materials and Methods). Isotype scores ranged between -14.5 and 155.9, with a median of 105.9 (Figure 5; Supplementary Table S7). As observed for the 20 Thermococcaceae genomes, the distribution of isotype scores for the tRNAs encoded in the 210 archaeal genomes showed a large Gaussian-like peak near 110 and a long tail of low-scoring putative tRNA genes. tRNAscan-SE predicted 839 putative tRNA genes scoring below 85, accounting for ∼8 % of all predicted tRNA genes, and ∼94 % of all predicted possible pseudogenes. Notably, tRNAscan-SE predicts 742 standard tRNA genes with isotype scores below 85. On closer inspection, many of these are unlikely to be canonically functional (*e*.*g*., are predicted to encode truncated tRNAs, or tRNAs with unusual secondary structures). Yet some, particularly those with isotype scores between ∼60 and ∼90, are predicted to form tRNAs that may indeed be functional (for an example, see *Thermoplasmatales archaeon* BRNA1 Ala-CGC in Supplementary Text S1). We also examined the value of other types of scores generated by tRNAscan-SE (*e*.*g*., HMM score, infernal score) in identifying tRNA gene fragment but, compared to the isotype score, these resulted in a less clear division of canonical and non-canonical tRNA genes (see Supplementary Figure S2). In summary, while both the automated pseudogene detection and isotype scores are clearly informative, they currently do not offer a watertight separation of canonical and non-canonical tRNA genes in Archaea.

**Figure 5.**
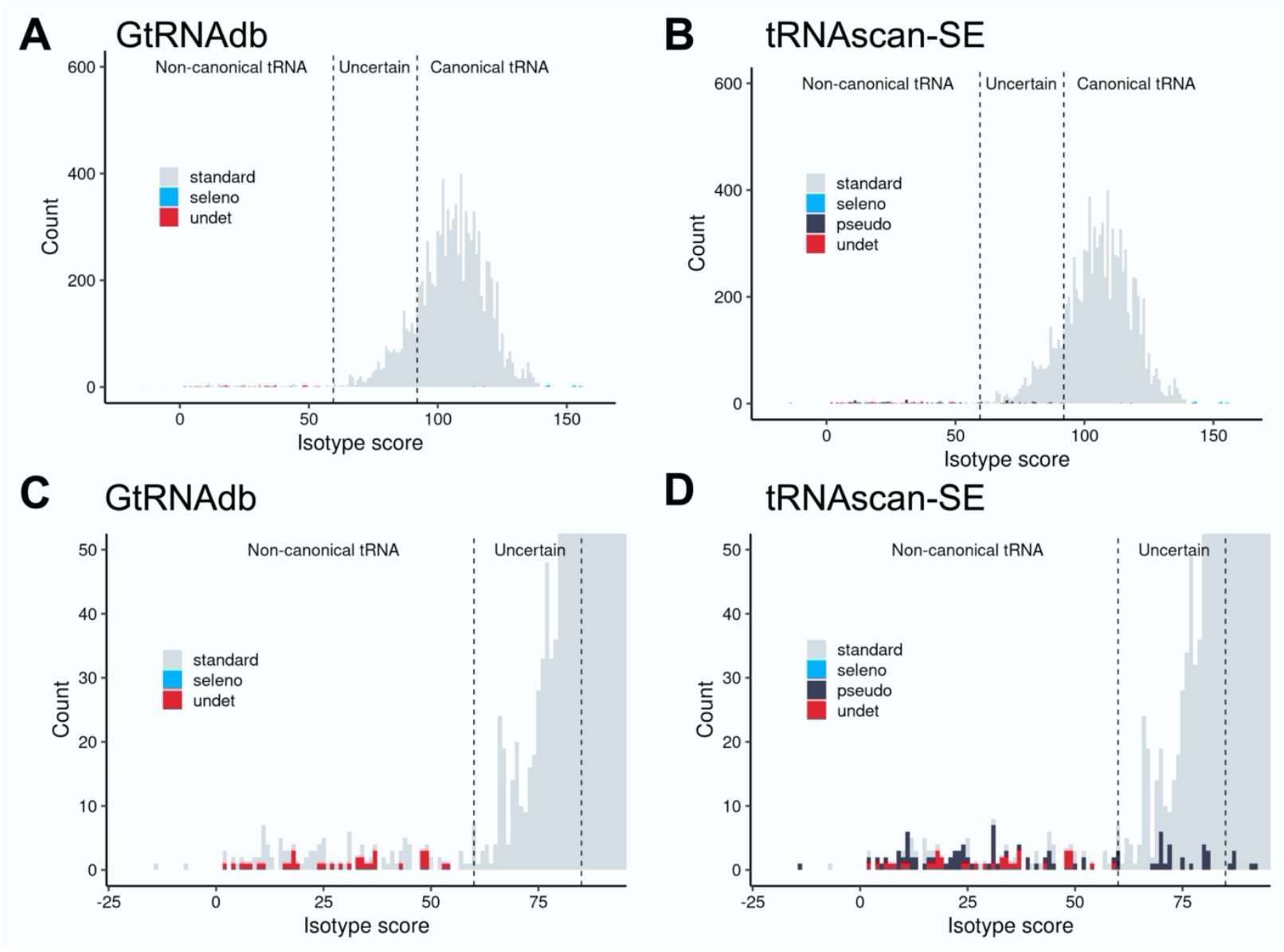
tRNAscan-SE isotype score as a predictor of canonical versus non-canonical tRNA genes in 210 archaeal genomes. (**A**) Histogram of the frequency of tRNA isotype scores from ∼10,000 tRNA genes from 210 archaeal genomes listed in (**A**) GtRNAdb, and (**B**) predicted by locally run tRNAscan-SE. (**C-D**) Closeup of isotype scores 0-100 in panels A and B. Note that in panels A and C (GtRNAdb) the pseudogene category (‘pseudo’) is again absent, due to disabling of pseudogene detection function (see ‘Run Options/Stats’ tab in GtRNAdb listings). Category ‘seleno’ denotes putative selenocysteine tRNA genes. Dotted lines demonstrate the use of tRNAscan-SE isotype scores as a useful indicator of the nature of putative archaeal tRNA genes: <60=likely non-canonical, 60-90=uncertain (*i*.*e*., may be either canonical or non-canonical), >90=likely canonical.

## DISCUSSION

In this work, we have performed a detailed manual analysis of the computationally predicted tRNA gene sets listed in GtRNAdb for each of 20 high quality genome sequences from the archaeal Thermococcaceae family. Foremost, we highlight that the tRNAscan-SE software has done a truly impressive job of identifying putative tRNA genes in these organisms; with a degree of manual curation, sensible and translationally complete canonical tRNA gene sets are recognizable in all 20 organisms (see Figure 1B). Additionally, and arguably equally usefully, tRNAscan-SE identifies a number of non-canonical tRNA gene fragments resulting from biologically relevant and widespread phenomena: CRISPR-Cas activity and the integration of incoming genetic elements (see Table 1). However, the process of distinguishing between these two categories (canonical and non-canonical), particularly on the widely-used, public database GtRNAdb, leaves some room for improvement.

Our analyses provide insights that allow progress towards three goals: (i) obtaining estimates of the accuracy with which tRNAscan-SE predicts, annotates, and categorizes putative tRNA genes, (ii) identifying specific sources of error, with the aim of highlighting specific avenues for possible further improvement of tRNAscan-SE and/or interpretation of the output (both by GtRNAdb and independent users), and (iii) gaining insight into the properties and evolutionary dynamics of archaeal tRNA gene sets. Each of these is discussed below.

### (i) The accuracy of automated tRNA gene prediction, annotation, and categorization on GtRNAdb

In eleven independent cases (see Table 1, rows 1-11), our manual curation process yielded conclusively different annotations of predicted tRNA genes than those obtained by the automated process (listed in GtRNAdb). These included the complete removal of eight predicted tRNA genes from the canonical tRNA gene sets, and the re-annotation of a further three to other – previously absent – isotypes (Table 1, rows 9-11). Overall, fourteen apparent deviations were removed from a set of 929 predicted tRNA genes, yielding an error rate of ∼1.5 % for the 20 Thermococcaceae GtRNAdb listings.

### (ii) Observed sources of error in automated tRNA gene annotations, and possible solutions

When predicting the above tRNA gene sets, one source of error was the annotation of tRNA-like sequences as standard tRNA genes (or, less commonly, tRNA genes of unknown or unidentified isotypes) in GtRNAdb. Eight such instances were detected within our dataset of 20 Thermococcaceae genomes (see Table 1). The existence of these sequences was attributed to two distinct mechanisms: the integration of horizontally transferred genetic elements into host tRNA genes (seven instances), and CRISPR-Cas activity (one instance).

The first mechanism – integration of genetic elements – is a general process in both Archaea and Bacteria ^52,53^ and, as such, similar tRNA gene-like remnants are expected to be a pervasive feature of prokaryotic genomes. Indeed, in addition to the seven examples detected in our Thermococcaceae dataset, we manually identified numerous tRNA gene fragments – many of which are mis-categorized on GtRNAdb as canonical tRNA genes – resulting from phage integration events in Archaea and Bacteria (see Supplementary Table S3). For example, GtRNAdb lists *Pseudomonas fluorescens* SBW25 as carrying two copies of the Cys-GCA gene, with these occurring on either side of a putatively identified ∼121 kb genomic island ^61^. One copy is low-scoring (tRNA-Cys-GCA-2-1, isotype score 29.0), and the corresponding putative tRNA was not detected in the mature tRNA pool of this bacterium ^32^. Notably, a third possible source of genomic sequences with tRNA-like features in Bacteria is transfer-<u>m</u>essenger RNA (tmRNA) genes ^62^. Such tmRNA genes encode RNA products of a few hundred base pairs, which perform a role in the quality control of translation (as opposed to elongation). Since tmRNA molecules carry some tRNA features, they have the potential to be identified and annotated by tRNAscan-SE. An example is seen in the GtRNAdb listing for *Thiobacillus denitrificans* ATCC 25259: tRNA-Und-NNN-3-1 (91-bp; isotype score 23.7) is actually the 3’ end of a tmRNA gene.

The accurate detection and annotation of pervasive tRNA-like genomic sequences poses a clear challenge to any tRNA identification software. To minimize the resulting overestimates of canonical tRNA genes in Archaea and Bacteria, we recommend that users predict canonical tRNA gene sets – or indeed, non-canonical tRNA gene sets – of interest using the following process: (i) pre-screening the genomes of interest for integrative genetic elements, tmRNA genes, and CRISPR loci, and flagging any partial and full tRNA sequences associated with such regions (numerous computational databases and prediction tools are available for the identification of such elements ^8,55,63–69^), (ii) using locally run tRNAscan-SE, paying particular attention to the pseudogene detection settings, and (iii) manually inspecting tRNAscan-SE output, using the tRNAscan-SE isotype score to focus on tRNA genes with isotype scores below ∼85 (many of which are likely to encode non-canonical tRNA genes), and any possible pseudogenes detected. Further, we suggest that the addition of two new tRNA gene categories to the output of tRNAscan-SE – *attR* (in line with established nomenclature denoting the ends of integrative elements; *e*.*g*., ^54^), and tmRNA – would be both useful and biologically meaningful.

A second source of error in our dataset was the presence of N nucleotides in input genome sequence leading to the mis-annotation of putative tRNA genes. Every such mis-annotation generated two errors: a false positive tRNA gene (due to the erroneous presence of the mis-annotated tRNA gene), and the corresponding false negative (due to the erroneous absence of the correctly annotated tRNA gene). Three examples were detected in our dataset, resulting in six errors (Table 1). This source of error could be minimized by a combination of users excluding (or, at a minimum, noting) genomes containing significant numbers of Ns, and/or by the addition of a note to the tRNAscan-SE output when Ns are detected within a putative tRNA gene sequence.

### (iii) The properties and evolutionary dynamics of archaeal tRNA gene sets

The results reported here provide insight into the evolutionary dynamics of tRNA gene sets in Archaea. Primarily, while the GC content of the 20 analyzed genomes varies widely (between 40 % and 56 %; see Supplementary Table S1), the predicted tRNA gene sets exhibit remarkable stability; with the likely exception of a single duplicate tRNA gene in *Pa. pacificus* DY20341, all 20 organisms contain a tRNA gene set encoded by 46 single-copy tRNA genes (see Figure 1B). These observations are consistent with previous suggestions of a ‘standard’ archaeal tRNA gene set ^4,5^, with occasional, small – and perhaps evolutionarily fleeting – instances of divergence.

## DATA AVAILABILITY STATEMENT

Raw data relating to this project is available at https://zenodo.org/record/6782366#.YsQSaZDP01A The extended TIGRFAM database can be found at https://zenodo.org/record/3839790#.YjByaVzMI3g The perl script used during phylogenetic tree construction, catfasta2phyml.pl, is available in the GitHub repository (https://github.com/nylander/catfasta2phyml) tRNAscan-SE is a freely available resource available online (http://lowelab.ucsc.edu/tRNAscan-SE/) GtRNAdb is a publicly accessible resource available online (http://gtrnadb.ucsc.edu/)

## ACKNOWLEDGEMENTS

None.

## SUPPLEMENTARY MATERIALS LIST AND LEGENDS

**Supplementary Figure S1. Phylogenetic tree highlighting the evolutionary relationships of the 20 Thermococcaceae genomes listed in GtRNAdb**. The complete genome sequences of all genomes were obtained (see Supplementary Tables S1, S6, and S7). Alignments were generated using 12,403 amino acid positions across 43 shared marker proteins (see Supplementary Table S5). The support shown next to the branches is based on 100 bootstrap replicates. Three genomes - *Methanococcus vannielii SB, Methanococcus maripaludis*, and *Methanococcus aeolicus* – were used to root the tree.

**Supplementary Figure S2. tRNAscan-SE isotype score can be used as an indicator of whether a putative archaeal tRNA gene is canonical or non-canonical**. Frequencies of locally run tRNAscan-SE (version 2.0.6)-derived scores for ∼10,000 putative tRNA genes in 210 archaeal genomes. (**A**) Isotype score (also presented in Figure 5B). (**B**) Infernal score, (**C**) HMM score, and (**D**) Secondary structure-only score. See Chan and Lowe 2021 (doi:10.1093/nar/gkab688) and references therein for a detailed description of score calculations.

**Supplementary Text S1. Primary and predicted secondary structures of putative tRNA genes of interest**. Primary sequences were obtained from GtRNAdb. Secondary structures and listed scores were predicted using the web based tRNAscan-SE 2.0 (sequence source: archaeal; search mode: default).

**Supplementary Table S1: Genomic characteristics of the 20 Thermococcaceae organisms**.

**Supplementary Table S2: Evidence for partial tRNA sequences resulting from the integration of genetic elements in the 20 *Thermococcaceae* genomes**. Genomic details of seven partial tRNA fragments resulting from the integration of prophages (*i*.*e*., erroneously identified as canonical tRNA genes; category 1b in Table 1) in *Thermococcaceae*.

**Supplementary Table S3: Evidence for partial tRNA sequences resulting from the integration of genetic elements across Archaea and Bacteria**. Genomic details of 17 partial tRNA fragments resulting from the integration of prophages in Archaea and Bacteria.

**Supplementary Table S4: Details of ribosomal RNA (*rrn*) operons and associated tRNA genes in the 20 *Thermococcaceae* genomes**. Only one genome differs from the norm of a single *rrn* operon and Ala-TGC gene: *Pa. pacificus* DY20341 carries two *rrn* operons and thus two Ala-TGC genes (*i*.*e*., a real deviation; category 3).

**Supplementary Table S5: Details of markers for drawing of the Thermococcaceae phylogenetic tree**.

**Supplementary Table S6: tRNA gene predictions, annotations, and scores for 20 Thermococcaceae organisms**. Tab 1: downloaded from GtRNAdb. Tab 2: predicted using locally run tRNAscan-SE (version 2.0.6, as for the GtRNAdb entries in Tab 1).

**Supplementary Table S7: tRNA gene predictions, annotations, and scores for 210 Archaea**. Tab 1: downloaded from GtRNAdb. Tab 2: predicted using locally run tRNAscan-SE (version 2.0.6, as for the GtRNAdb entries in Tab 1).

