## Supplementary Figure S1 for "On distinguishing between canonical tRNA genes and tRNA gene fragments in prokaryotes"

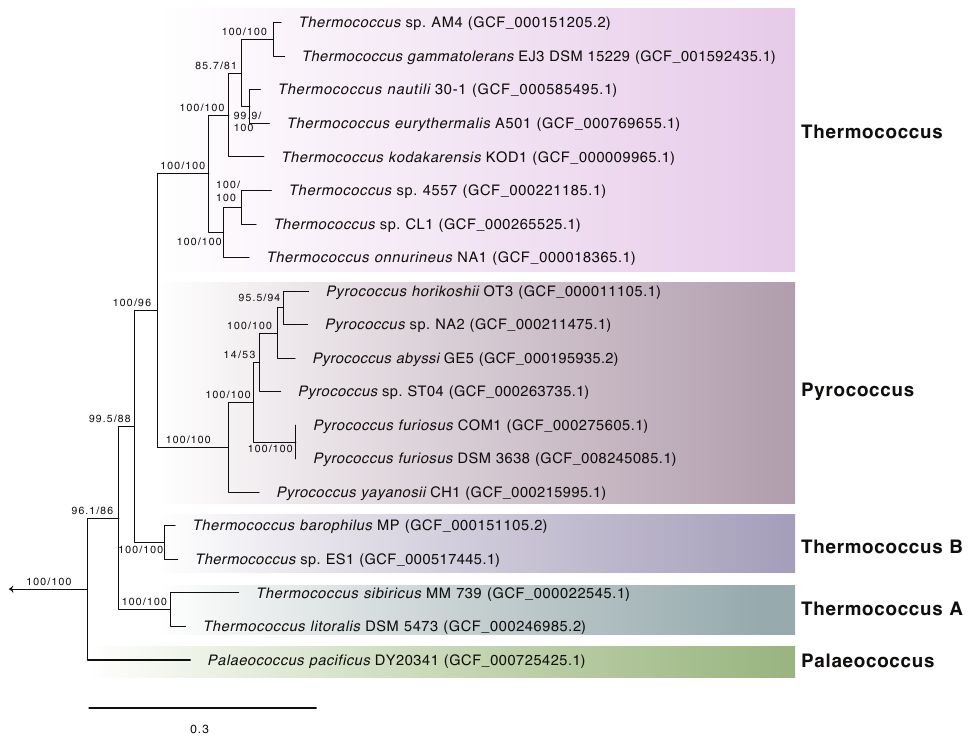

**Supplementary Figure S1. Phylogenetic tree highlighting the evolutionary relationships of the 20 Thermococcaceae genomes listed in GtRNAdb.** The complete sequences of all genomes were obtained (see Supplementary Tables S1 and S5). Alignments were generated using 12,403 amino acid positions across 43 shared marker proteins (see Supplementary Table S5). A maximum-likelihood phylogenetic tree was inferred with the LG+C60+F+R model with a SH-like approximate likelihood tests (left) and ultrafast bootstrap approximation (right), each run with 1000 replicates. Three genomes – *Methanococcus vannielii SB*, *Methanococcus maripaludis*, and *Methanococcus aeolicus* – were used to root the tree. See Materials and Methods for further details.
