## Supplementary Figure S2 for "On distinguishing between canonical tRNA genes and tRNA gene fragments in prokaryotes"

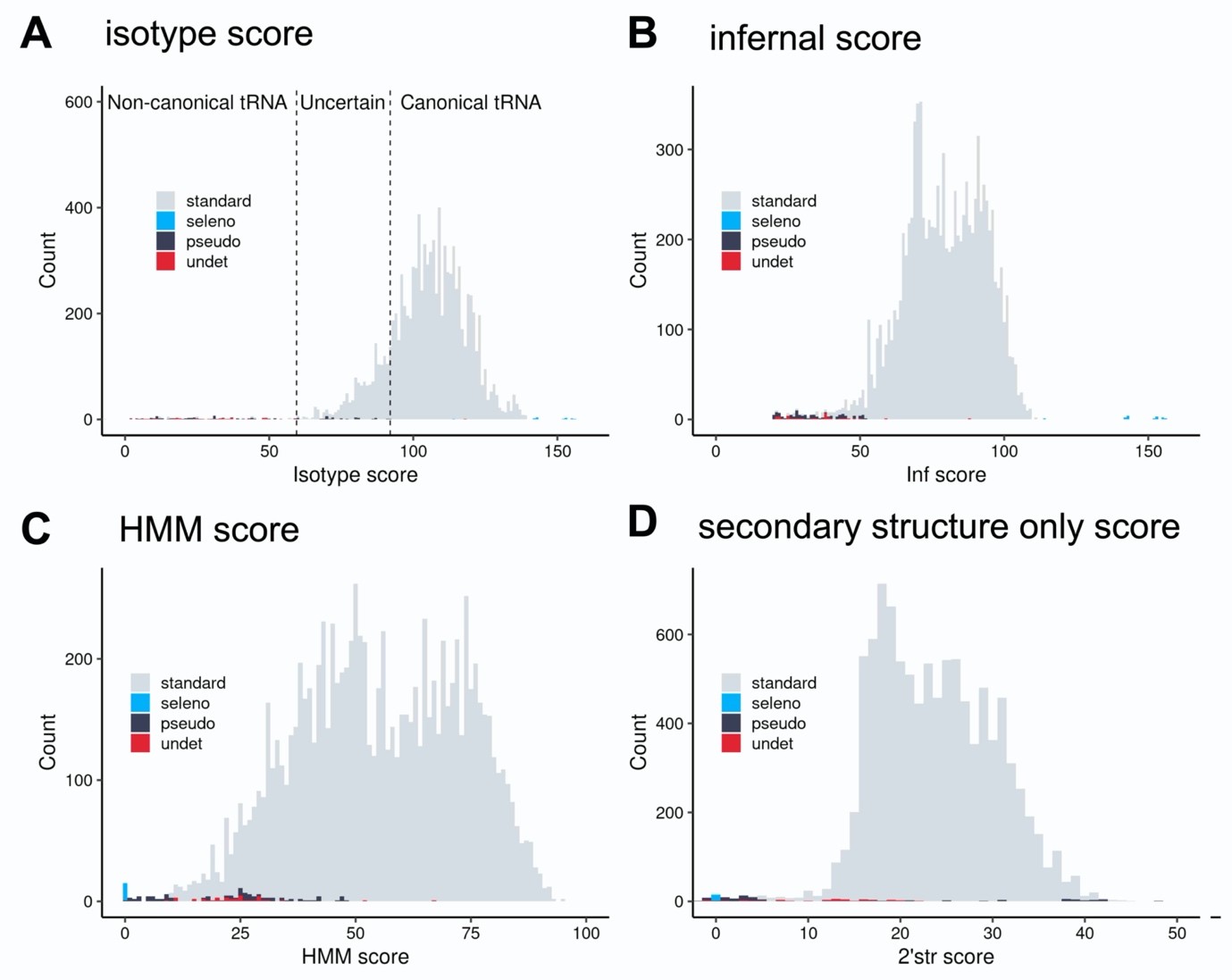

**Supplementary Figure S2. tRNAscan-SE isotype score can be used as an indicator of whether a putative archaeal tRNA gene is canonical or non-canonical.** Frequencies of locally run tRNAscan-SE (version 2.0.6)-derived scores for ~10,000 putative tRNA genes in 210 archaeal genomes. (**A**) Isotype score (also presented in Figure 5B). (**B**) Infernal score, (**C**) HMM score, and (**D**) Secondary structure-only score. See Chan and Lowe 2021 (doi:10.1093/nar/gkab688) and references therein for a detailed description of score calculations.
