## Supplementary Text S1 for "On distinguishing between canonical tRNA genes and tRNA gene fragments in prokaryotes"

**Primary and predicted secondary structures of putative tRNA genes of interest**

*1	Ser-CGA (+) in T. gammatolerans EJ3*

>T_gammatolerans_Ser-CGA-2-1

GGAGTAGCCTTCTAAGCCGGAGGtCGCGGGTTCGAATCCCGCCGGGCCCGCCA

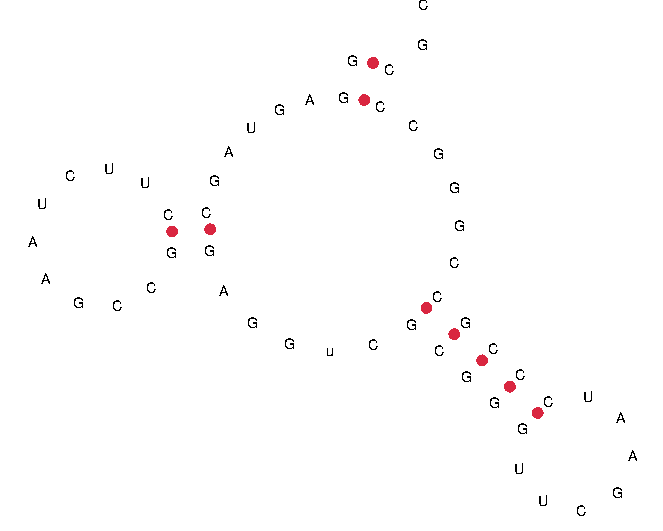

Infernal Score: 22.6

Isotype Score:
10.4

*2	Leu-CAA (+) in T. nautili 30-1*

>T_nautili_Leu-CAA-2-1

GGAGCGGTAGCGGTaACCCCAGCTTTCAAttctcctcgagtcttattgcaacttcctccttctgcaggccgataccgcaccccttatcctttcaattctcctcgagtcttattgcaacacgctcgaacagcacataaaagcgatagctgacgcctttcaattctcctcgagtcttattgcAACGTTGGtCGGGGGTTCAAATCCCCTCCccgGCTCCA

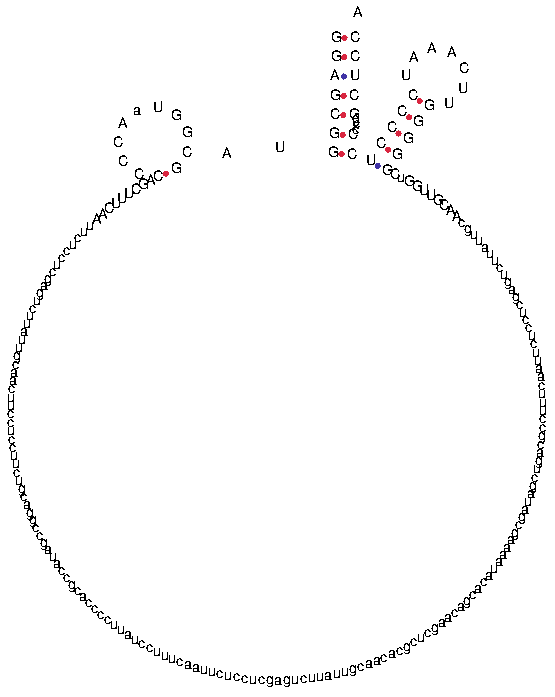

Infernal Score: 22.2

Isotype Score:
2.3

*3	Arg-TCT (+) in T. kodakarensis KOD1*

>T_kodakarensis_Arg-TCT-2-1

GGCCCTTTCAGCAGCccacaattcattaAGGGAACGGCCTTCTAAGCCGGAGGtCGCGGGTTCGAATCCCGCCGGGCCCGCCA

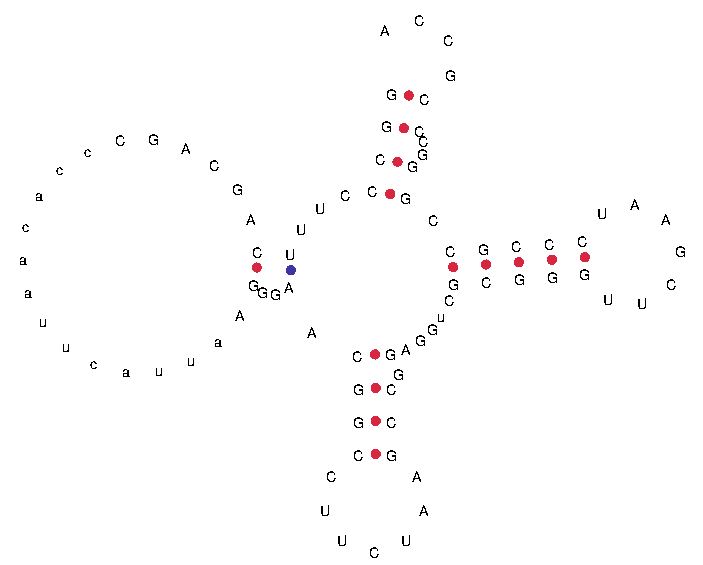

Infernal Score: 27.1

Isotype Score:
21.4

*4	Val-CAC (+) in T. kodakarensis KOD1*

>T_kodakarensis_Val-CAC-2-1

GGTGTCCcAACAGTCGTTtAGACTGCCCTCACACGGCGGAGGtCCGGGGTTCGAATCCCCGCGGGCCCACCA

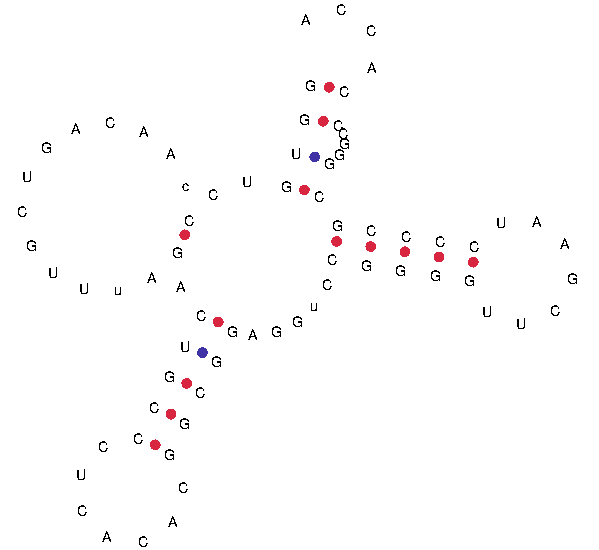

Infernal Score: 45.3

Isotype Score:
83.4

*5	Gln-TTG (+) in P. yayanosii CH1*

>P_yayanosii_Gln-TTG-2-1

AGCCCTGTAagaCCTTCATTtGGCgaatAAAGGcttttccgACGGGCTTTGGATCCCGCGACTCGGGTTCGAATCCCGGCGGGGCTACCA

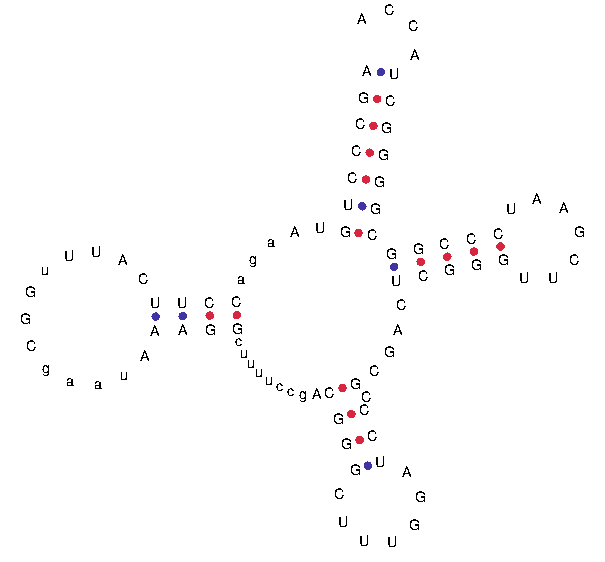

Infernal Score: 26.5

Isotype Score:
13.4

*6	Ser-CGA (+) in T. sp ES1*

>T_ES1_Ser-CGA-2-1

GGCTGTTTTGCGGTttTGCTGaCCCTTACGAGGCGGAGGtCCGGGGTTCGAATCCCCGCGGGCCCACCA

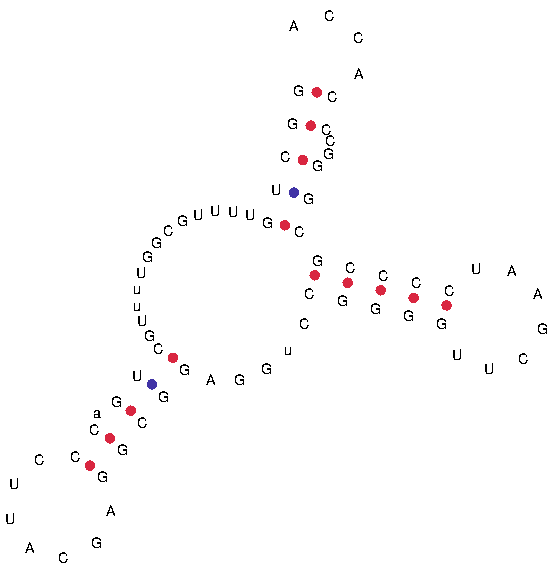

Infernal Score: 33.6

Isotype Score:
30.9

*7	Und-NNN (+) in T. sp ES1*

>T_ES1_Und-NNN-1-1

GGGTTTGTAGAGGGgAACAAGCCTTCTGctgggtctattagctcaatagccgatgacaaactctctaaattctccaaaaaccgctttcgAGCCCGCGcCCCGGGTTCAAATCCCGGCCGGGGCACCA

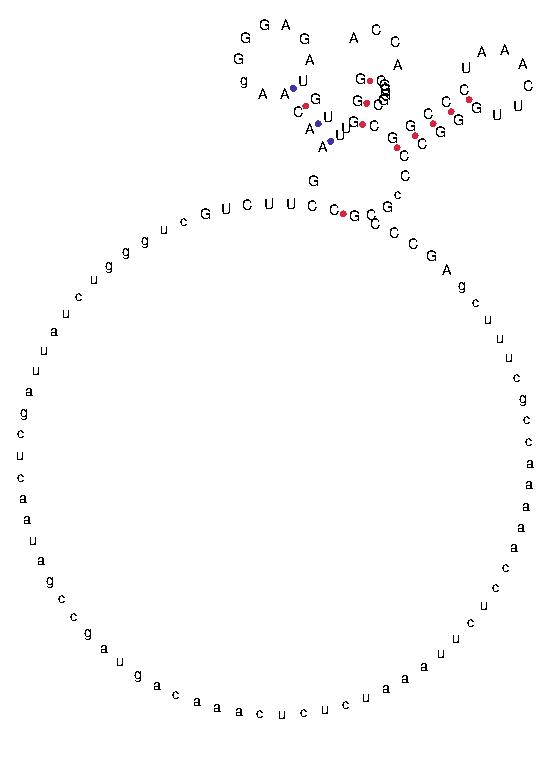

Infernal Score: 25.5

Isotype Score:
16.8

*8	Leu-CAA (+) in T. litoralis DSM 5473*

>T_litoralis_Leu-CAA-2-1

GGGTCGTTTTTtGGATtTAATagtttagacggctgtgGATCCCCTAGCCCGGGTTCAAATCCCGGCCCCGGCCCCA

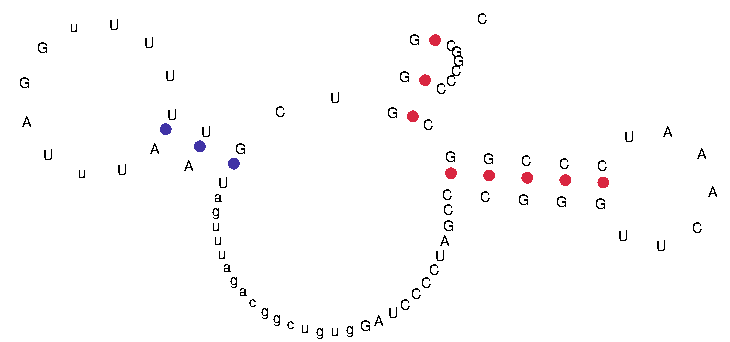

*9 & 10	Leu-NAG (+) and Leu-CAG (-) in T. litoralis DSM 5473*

>T_litoralis_Leu-NAG-1-1 (Leu-CAG)

GCGGGGGTTGCCGAGCCtGGTcaAAGGCGCGGGATTNAGGGTCCCGTCCCGTAGGGGTtCCGGGGTTCAAATCCCCGCCCCCGCACCA

**Leu-CAG**
(suggested)

**Leu-NAG**

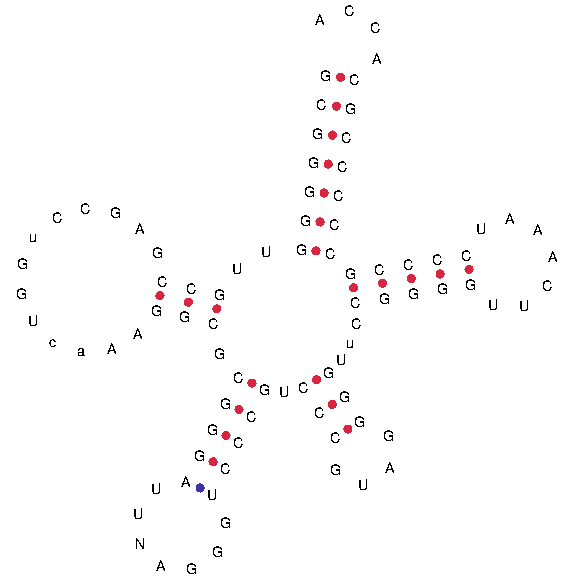

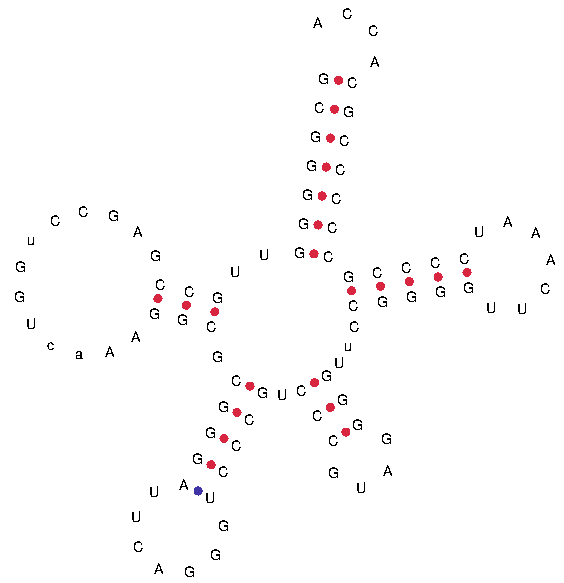

Isotype Score:
134.9

Isotype Score:
135.1

Infernal Score: 94.2

Infernal Score: 93.6

*11 & 12	Pro-NGG (+) and Pro-GGG (-) in T. litoralis DSM 5473*

>T_litoralis_Pro-NGG-1-1 (Pro-GGG)

GGGGCCGTGGGGTAGCTtGGTctATCCTNCCGGCTTNGGGNGCCGGAGACCCGGGTTCAAATCCCGGCGGCCCCACCA

**Pro-GGG**
(suggested)

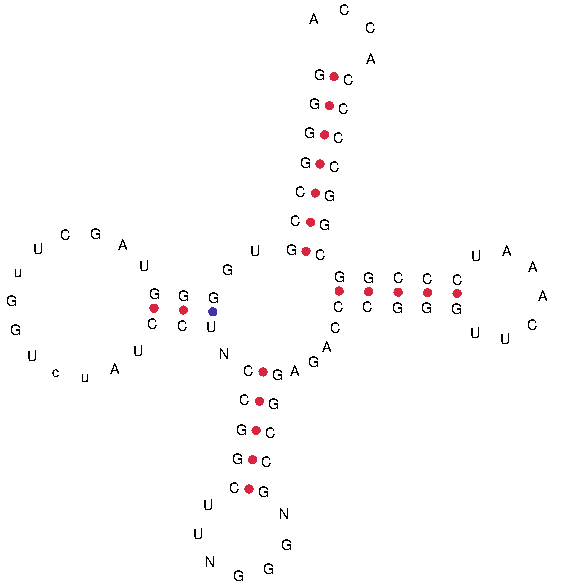

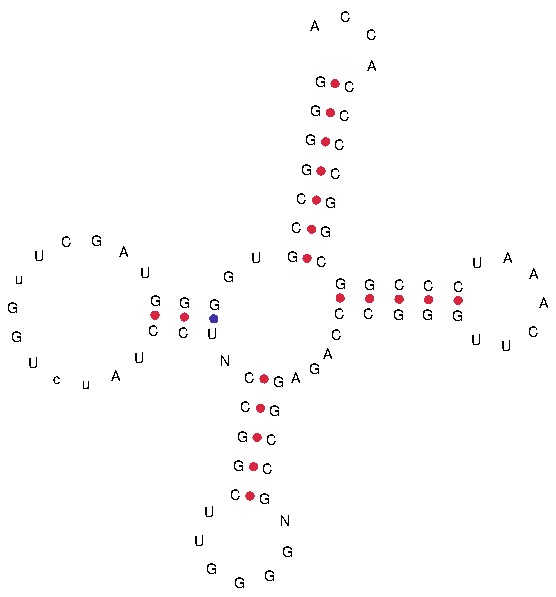

Isotype Score:
120.3

Isotype Score:
120.3

**Pro-NGG**

Infernal Score: 91.4

Infernal Score: 91.4

*13 & 14	Und-NTG (+) and Gln-CTG (-) in T. litoralis DSM 5473*

>T_litoralis_Und-NTG-1-1 (Gln-CTG)

AGCCCCGTGGTGTAGCGGCcaAGCATGCGGGACTNTGGATCCCGCGACCGGGGTTCGAATCCCCGCGGGGCTACCA

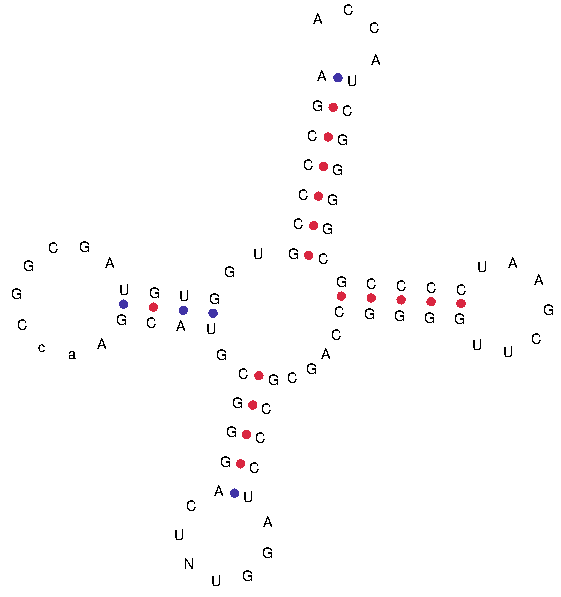

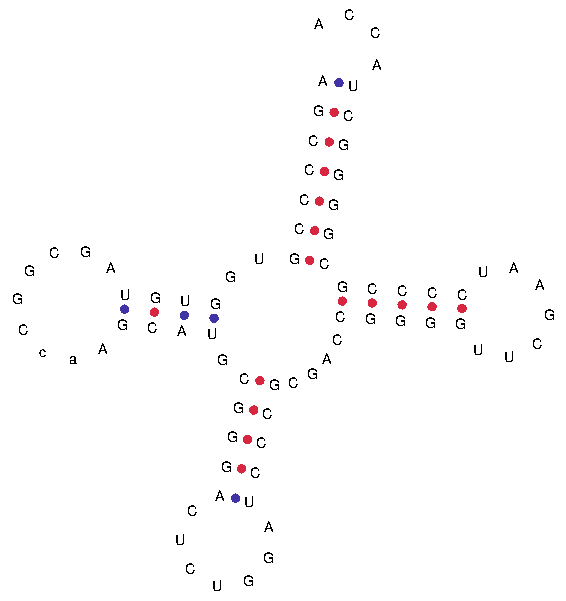

Isotype Score:
117.9

Isotype Score:
117.9

Infernal Score: 89.1

**Gln-CTG**
(suggested)

**Und-NTG**

Infernal Score: 88.4

*15	Ala-TGC (+) in Pa. pacificus DY20341*

>Pa_pacificus_Ala-TGC-1-2

GGGCCGGTAGCTCAGCCtGGGAGAGCGCCGGCTTTGCAAGCCGGAGGcCCCGGGTTCAAATCCCGGCCGGTCCACCA

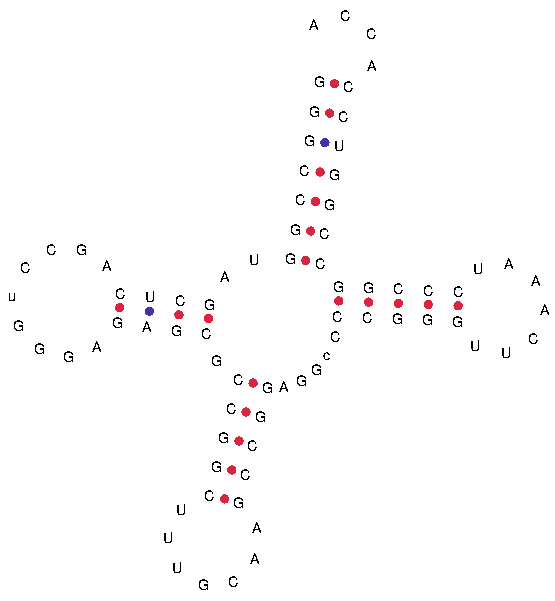

Infernal Score: 102.2

Isotype Score:

122.6

*Figure 4: Predictions for the* P. furiosus *Arg-GCG tRNA genes*

In Figure 4C-D, the Arg-GCG tRNA gene is from *P. furiosus* DSM3638; it appears to be a true standard tRNA gene with a relatively low tRNA Model Score (85.5). This relatively low score seems to result from a missing ‘G’ in the later part of the primary sequence, generating a short acceptor stem with an unpaired ‘C’. A close relative, *P. furiosus* COM1*,* carries a Arg-GCG containing the missing ‘G’ base. Hence, we propose this is either an error in the genome sequence of *P. furiosus* DSM3638 genome sequence (or, possibly, a relatively recent mutation).

>1_Pfuriosus_DSM3638_Arg-GCG-1-1

GCCCCGGTGGCCTAGCCtGGAtAGGGCGCGAGGCTGCGGACCTCGAGGtCCGGGGTTCAAATCCCCGCCGGGCGCCA

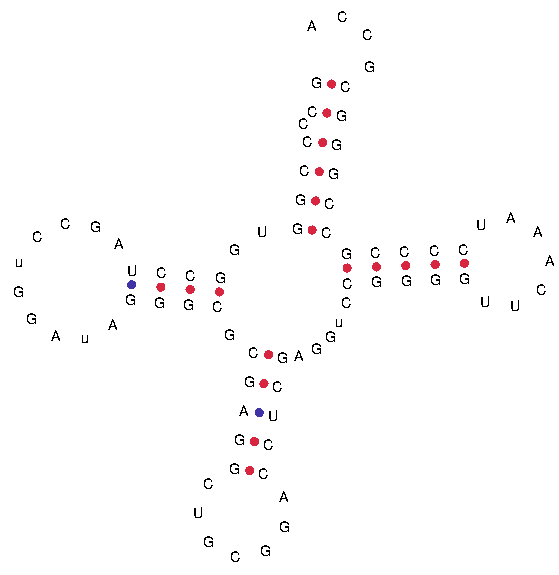

Infernal Score

85.5

Isotype Model Score

88.1

Unpaired base

>2_Pfuriosus_COM1_Arg-GCG-1-1

GCCCCGGTGGCCTAGCCtGGAtAGGGCGCGAGGCTGCGGACCTCGAGGtCCGGGGTTCAAATCCCCGCCGGGGCGCCA

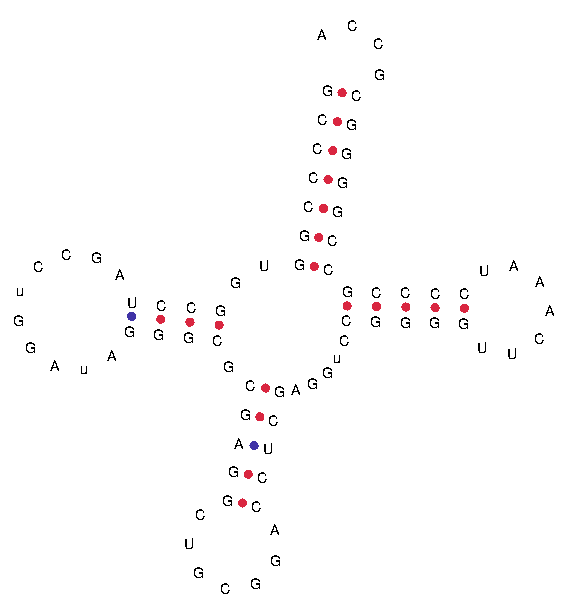

Infernal Score

101.0

Isotype Model Score

102.5

Paired bases

*Low-scoring tRNA gene example: Ala-CGC from* Thermoplasmatales archaeon *BRNA1*

The GenBank accession for this genome is CP002916.1. We note that it is predicted by tRNAscan-SE to encode a relatively low-scoring (isotype score) tRNA gene that could conceivably encode a functional, canonical tRNA:

tRNA-Ala-CGC-1-1

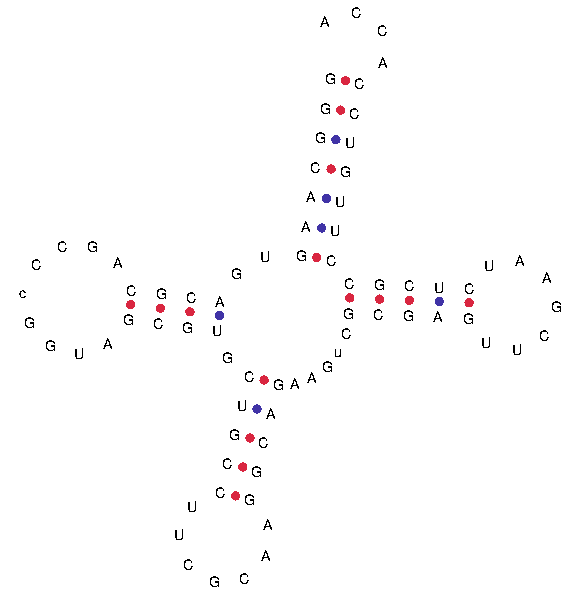

Infernal Score

79.8

Isotype Model Score

83.7
